# Wild pollinator activities negatively related to honey bee colony densities in urban context

**DOI:** 10.1101/667725

**Authors:** Lise Ropars, Isabelle Dajoz, Colin Fontaine, Audrey Muratet, Benoît Geslin

## Abstract

As pollinator decline is increasingly reported in natural and agricultural environments, cities are perceived as shelters for pollinators because of low pesticide exposure and high floral diversity throughout the year. This has led to the development of environmental policies supporting pollinators in urban areas. However, policies are often restricted to the promotion of honey bee colony installations, which resulted in a strong increase in apiary numbers in cities. Recently, competition for floral resources between wild pollinators and honey bees has been highlighted in semi-natural contexts, but whether urban beekeeping could impact wild pollinators remains unknown. Here, we show that in the city of Paris (France), wild pollinator visitation rates is negatively correlated to honey bee colony densities present in the surrounding (500m – slope = −0.614; p = 0.001 – and 1000m – slope = −0.489; p = 0.005). More particularly, large solitary bees and beetles were significantly affected at 500m (respectively slope = −0.425, p = 0.007 and slope = - 0.671, p = 0.002) and bumblebees were significantly affected at 1000m (slope = - 0.451, p = 0.012). Further, lower interaction evenness in plant-pollinator networks was observed with honey bee colony densities within 1000 meter buffers (slope = −0.487, p = 0.008). Finally, honey bees tended to focus their foraging activity on managed rather than spontaneous plant species (student t-test, p = 0.001) whereas wild pollinators equally visited managed and spontaneous species. We advocate responsible practices mitigating the introduction of high density of hives in urban environments. Future studies are needed to deepen our knowledge about the potential negative interactions between wild and domesticated pollinators.

## Introduction

The recent decline of pollinating insect populations is driven by a conjunction of factors, including habitat fragmentation, use of pesticides, multiplication of pathogens, global warming and the decline of the wild flora [1]. Agricultural landscapes have changed, harbouring fewer floral resources and habitats to support diverse pollinating communities [2,3]. Consequently, many agricultural landscapes are becoming less conducive for pollinators and for beekeeping activities [4]. At the same time, areas that were previously rarely exploited by beekeepers are now under strong pressure to receive apiaries; this is the case in natural habitats and cities [5,6]. Indeed, cities harbour diverse plant species flourishing all year long due to management practices [7] and heat island effect, thus providing resources throughout the year for pollinators [8]. The low pesticide policies applied in many conurbations also may create favourable conditions for the maintenance of diverse pollinator communities [9]. In parallel, inhabitants have associated *A. mellifera* to the quality of their environment, and honey bees have become a symbol of biodiversity for media and general public [10]. Many citizens have thus installed colonies as their own contribution to mitigate the pollinator decline [11,12] and urban introductions of honey bee colonies have been promoted by public authorities and decision makers. In many cities, this has translated into very recent and rapid increases in the number of honey bee colonies (10 colonies per km^2^ in London – United Kingdom [13], 15 colonies per km^2^ in Brussels – Belgium [14]).

However, cities are not depauperate in wild pollinating insects and there is increasing evidence that they host diverse assemblages of wild bees [15,16]. This has led to rising concern about numerous introductions of honey bees in cities, that may negatively impact the wild pollinating fauna through competition for floral resources [11]. In other habitats, such as semi-natural (calcareous meadows [17] or scrubland [18,19]) or agricultural landscapes, several authors have detected exploitative competition between domesticated and wild pollinators through the monopolization of floral resources by honey bees [20,21]. However, it is largely unknown to what extent honey bee introductions in cities could impact wild pollinator communities and their foraging activity on urban plant communities. Moreover, the effect of increasing honey bee densities has rarely been assessed using network approaches [11]. Massively introduced honey bees might impair the pollination function at community level by, for example, focusing their visits on managed (ornamental) plant species rather than spontaneous ones [11]. Here, we explore those issues in the city of Paris (France), which has recently experienced a strong growth of its honey bee populations within a few years. In 2013, Paris hosted 300 honey bee colonies, and in 2015 this figure had more than doubled, reaching 687 colonies, corresponding to 6.5 colonies.km^-2^ (data of the veterinary services of Paris; Fig 1), and has continued to increase since. In this context, our first objective was to analyze the effect of increasing honey bee colony density on the visitation rates of wild pollinators at the community and morphological group levels. Secondly, we explored how the evenness of plant-pollinator networks was affected by increasing honey bee colony densities. Finally, we investigated the floral preferences of wild and domesticated pollinators for managed or spontaneous plant species.

**Fig 1.**
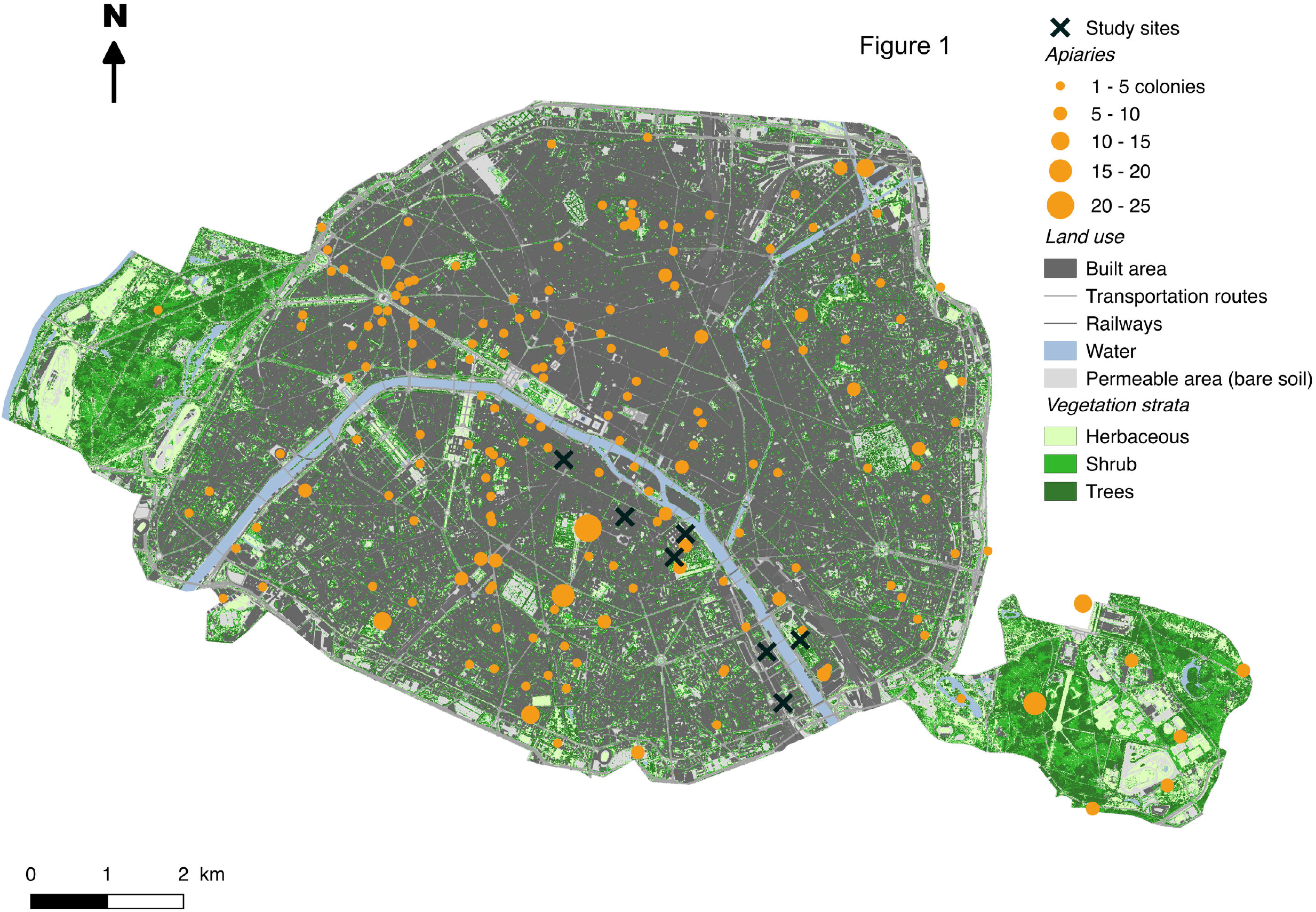
Localisation of honey bee colonies and study sites in the city of Paris. Vegetation height and land use map were obtained from APUR database (http://opendata.apur.org/datasets/).

## Methods

### Study sites and plant-pollinator survey

The city of Paris (48°51’12’’ N, 2°20’55’’ E, Île-de-France, France) is a densely populated urban area (2 220 445 inhabitants in 2014, 105km^2^). In this city, for three consecutive years, we monitored plant-pollinator interactions in five (in 2014) to seven (in 2015 and 2016) green spaces. We chose these green spaces by their contrasted densities of honey bee colonies in their surroundings (Figs 1 and 2) and for their relative accessibility (access granted by the Bibliothèque nationale de France, campus of Paris Diderot University, Pierre et Marie Curie University, Descartes University, and 3 gardens monitored by the Paris Direction des Espaces Verts et de l’Environnement). Sites were distant from 410 to 6 264 meters (see S2 Table). From May to July 2014 and from April to July 2015 and 2016, we carried respectively 8, 11 and 13 observation rounds per green space, spaced out at least by a week. For each round, we focused our observations on three one-meter squared patches chosen to be well-flourished. For each flower visit, we identified the visited plant to the lowest possible taxonomical level according to the taxonomic repository of France [22] and classified it as managed or spontaneous (see S1 Table). Mean richness of visited plant species within patches could vary from 2.5 to 6.5 species depending on the flowering phenology of the plants present in the site.

**Fig 2.**
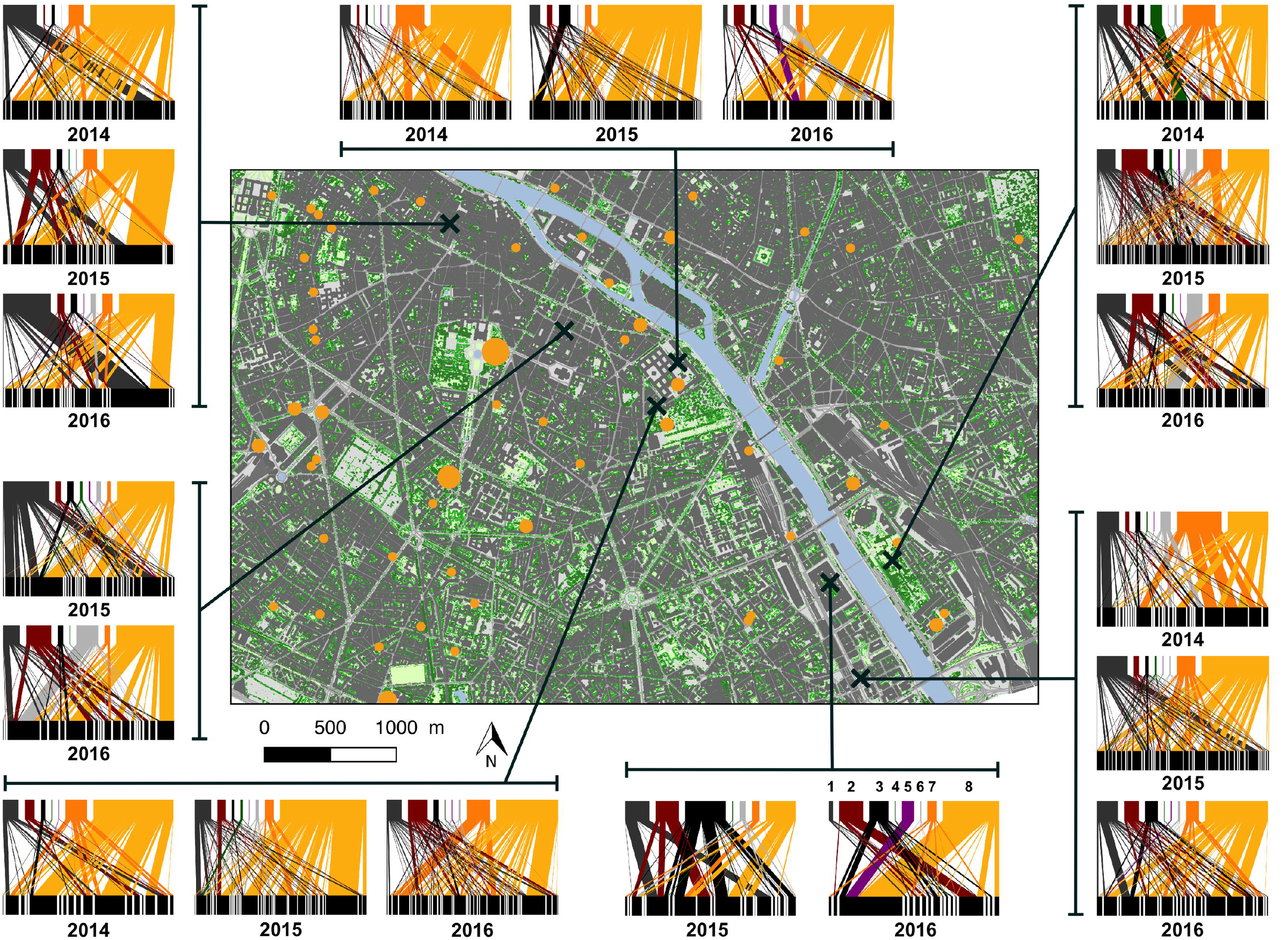
Study sites within Paris (France) and related plant-pollinator networks. Bipartite networks are retrieved from the compilation of all observed interactions between pollinators (top bar) and plants (bottom bar). Numbers under networks indicate observations year. Each pollinator block represents a morphological group and each plant block represents a species. The width of links is proportional to the number of interactions *(i.e.* pollinating activity). Pollinator morphological groups are classified by order (left to right) colours and numbers: dark grey-1, small solitary bees; dark red-2, large solitary bees; black-3, syrphids; dark green-4, beetles; purple-5, butterflies; flies-6, light grey; orange-7, bumblebees; yellow-8, honey bees. Vegetation height and land use map came from APUR database (http://opendata.apur.org/datasets/).

Further, we classified the insect visitors into one of eight morphological groups: small and large solitary bees, honey bees – *A. mellifera,* bumblebees, beetles, butterflies, hoverflies and other flies [23]. Observation rounds were performed during warm sunny days with no wind (<15°C) and were carried out between 9 a.m. and 7 p.m.

### Honey bee colonies location

The law requires beekeepers to report their honey bee colonies to the veterinary services of the city. This is to our knowledge the most accurate data existing to date regarding the location of honey bee colonies within Paris – even if we are aware that some beekeepers may not report their colonies. We used these data to estimate honey bee colony density within 500- and 1000-meter buffers centred on the study sites using the ArcGIS software (Version 10.2). We chose these buffer sizes to match the mean foraging distances of the majority of wild and domesticated bees species [24,25].

### Statistical analyses

#### Spatial auto-correlation analysis

All statistical analyses were performed using R 3.2.2 (R Development Core Team, 2015). We checked the absence of spatial autocorrelation between our sites and honey bee colonies. We generated a matrix of distances between sites (see S2 Table), and built matrices using the number of honey bee colonies in 500m and 1000m around our sites. Mantel tests were carried out between these matrices. No significant spatial autocorrelation was observed for all 2 buffer sizes (respectively, 500m – p = 0.749; 1000m – p = 0.204). The same spatial autocorrelation test was significant for the visitation rates of whole wild pollinators (p = 0.025) but was not for morphological groups taken separately. Therefore, we added an autoregressive process of order 1 correlation structure (addition of site coordinates and a random effect with years nested within sites) in models which contained wild pollinator visitation rates as the variable to explain.

#### Floral resources

Floral resources can affect the composition and activity of pollinator assemblages [26]. To account for the spatial heterogeneity of floral resources availability surrounding the study sites, we combined the area covered by the herbaceous, shrub and tree strata with an estimation of the average production of floral unit per stratum along the observation campaign from February to July. We used a map of vegetation height, with a 50cm^2^/pixel resolution provided by the Parisian Urbanism Agency (APUR – http://opendata.apur.org/datasets/hauteur-vegetation-2015) to calculate the area covered by vegetation of less than 1 meter high (herbaceous stratum), between 1 and 10 meters (shrub stratum), and higher than 10 meters (tree stratum – Fig 1), this within buffers of 500 and 1000 meters centred on our study sites using Geographic Information Systems (GIS, ESRI ARC INFO v. 10.0). To estimate the floral resource production for shrubs and trees along the 6 months, we multiplied their area by their number of floral units/m^2^. AgriLand database [27] (S3 Table) allowed us to estimate the number of floral units/meter^2^ at flowering peak. For these two strata, we considered that the flowering period lasted for 1 month. For the herbaceous stratum, considering the flowering phenology, we modelled a normal distribution pattern (μ = 3; σ^2^ = 1.220) for 6 months, with the peak of floral production (2700 floral units / m^2^) occurring in the 3^rd^ month. Using this method, we averaged the number of floral units at 1,371 per month for the 6 months of flowering. Although not targeting urban areas, AgriLand database is the most comprehensive database on floral unit production allowing to account for difference in floral resources production among vegetation stratum.

To assess the local floral resources, we also calculated the mean number of visited plant species by pollinators of all patches observed per year as an estimation of site’s species richness.

#### Foraging activity analysis

To standardize the observation effort among years, we calculated visitation rates as the number of visits per minute and per flower visitor group on each site and for each year. Visitation rates were analysed using linear mixed effects models (lme, package “nlme”, R Development Core Team, 2015) and log transformed to approach normality. For each morphological group, fixed effects were a) the honey bee colony densities at 500 or 1000 meters around our sites, b) the estimation of the floral resources available in buffers of the same radius and c) the mean plant species richness of each site. We included the year nested within sites as a random effect to account for temporal repetitions. We performed model simplification based on the Akaike Information Criterion (AIC) and chose the best fit model with ΔAIC > 2 (dredge, package “MuMIn”, R Development Core Team 2015 – S5 Table) [28]. All variables were scaled to make their estimated effects comparable.

#### Plant-pollinator network analysis

To determine the impact of honey bee colony density on the structure of plant-pollinator network, we constructed 19 quantified interaction networks linking flower visitor morphological groups excluding honey bees to plant species, one per site and year. Interaction frequencies were calculated as the number of visits per minute. The structure of the interaction networks was assessed by the interaction evenness using the “bipartite” package [29].Interaction evenness is bounded between 0 and 1, and derived from the Shannon index, *H*= p_ij_log_2_p_ij_/log_2_*F*, where *F* is the total number of plant–pollinator interactions in the matrix and p_ij_ is the proportion of those interactions involving plant *i* and pollinator *j* [30,31]. This index reflects how balanced is the interaction strength between plants and pollinators. It decreases as the network is dominated by few highly frequent interactions and increase when the number of interactions is uniformly distributed [32]. We analyzed the interaction evenness using the same statistical models than for the visitation rate analysis, fixed effects were a) the honey bee colony densities at 500 or 1000 meters around sites, b) the estimation of the floral resources available in a buffer of the same radius and c) the mean plant species richness of each sites. A model simplification based on the Δ_AIC_ > 2 was used (S6 Table) [28]. We included the year nested within sites as random effect to account for temporal repetitions.

#### Floral preferences analysis

To assess the pollinator floral preferences of both wild pollinators and honey bees, we summed their visitations on managed or spontaneous plant species (S1 Table). To consider the respective availability of both plant types, visitation rates of pollinator groups on managed or spontaneous plants species were weighted by the percentage of managed and spontaneous species recorded at each site and year. The number of each plant type sampled per year is available in the S4 Table. Floral preferences were tested using Student t-tests comparing the visitation rates of pollinator groups between managed and spontaneous plants.

## Results

### Effect of honey bee colony densities on wild pollinator visitation rate

Pollinators were monitored for a total of 3,120 minutes during which we recorded 795 individual plant-pollinator links, totalling 32,694 visits on plants (16% of small solitary bees, 10% of large solitary bees, 12% of bumblebees, 1% of beetles, 6% of hoverflies, 4% of flies, 1% of butterflies and 50% of honey bees). 687 honey bee colonies were declared in Paris in 2015, which equates to an average density of 6.5 colonies/km^2^. Visitation rates of wild pollinators were negatively related to the density of honey bee colonies at both spatial scales (Fig 3 and 4, and Table 1, 500m – slope = −0.614; p = 0.001 – and 1000m–slope = −0.489; p = 0.005). Large solitary bees performed significantly fewer visits when the density of honey bee colonies increased within 500 meter buffers around our observation sites (Fig 3, Table 1, slope = −0.425; p = 0.007). This trend was significant for beetles too (Fig 3, Table 1, slope = – 0. 671; p = 0.002). The visitation rate of bumblebees significantly decreased when the density of honey bee colonies increased within 1000 meter buffers (Fig 4, Table 1, slope = – 0.451; p = 0.012). The visitation rate of honey bees was positively correlated with the number of honey bee colonies within 500 meter buffers (Fig 3, Table 1 – slope = 0.501; p = 0.020). However, we did not record any significant increase in the visitation rate of honey bees with the increased density of hives within 1000 meter buffers. Finally, we did not find any effects of honey bee colony density on the visitation rate of other morphological groups of pollinators such as small solitary bees, flies, hoverflies and butterflies (Δ_AIC_ < 2 between null models and convenience models or models contain only resources or richness variables – see S5 and S6 Tables).

**Fig 3.**
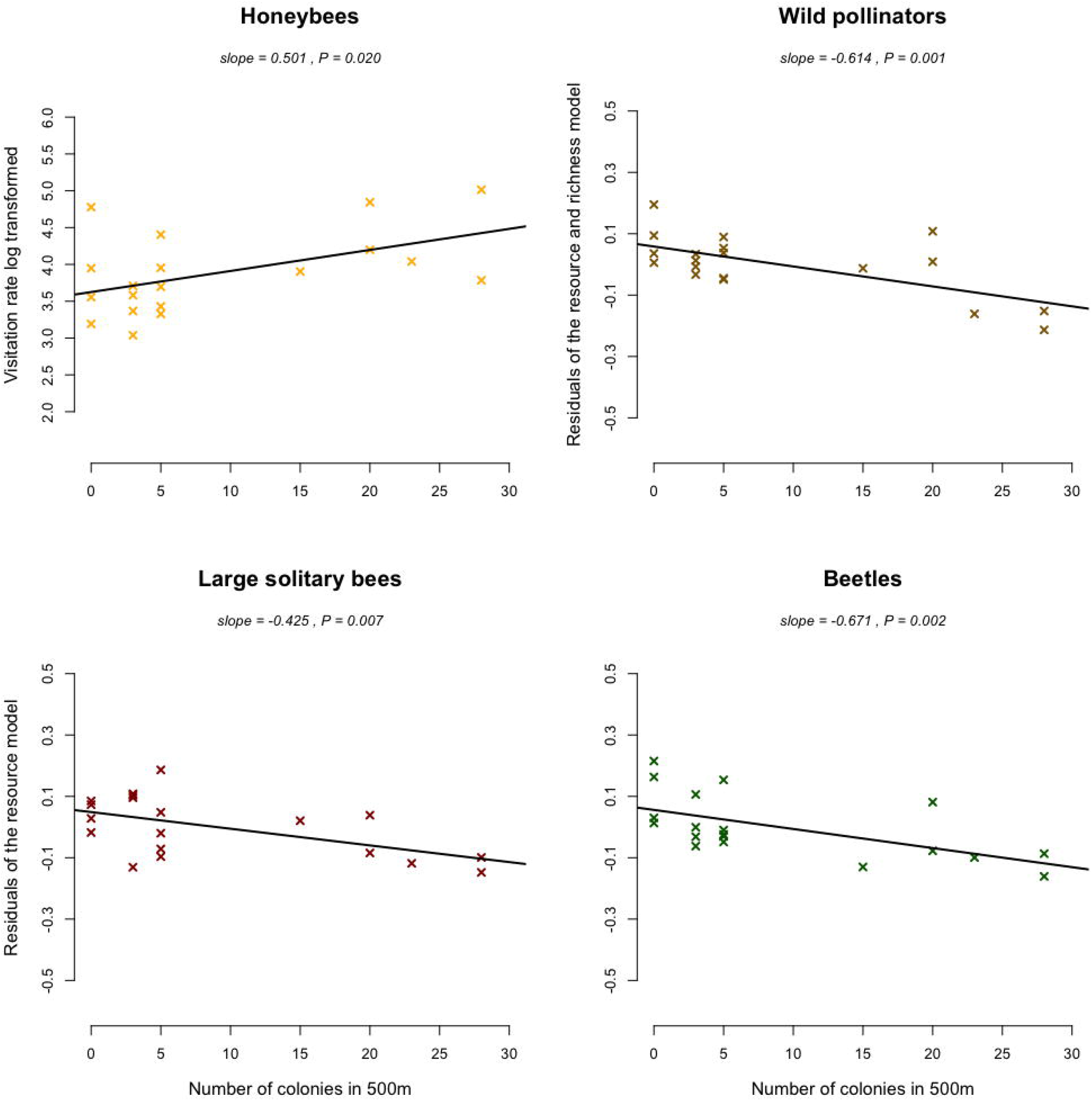
Morphological group visitation rates along the gradient of honey bee colony number in 500m around our observation sites. When the best model contained also the estimation of floral resources and/or richness as explanatory variable, residuals of morphological group visitation rates explained by resource variable only were used as y values.

**Fig 4.**
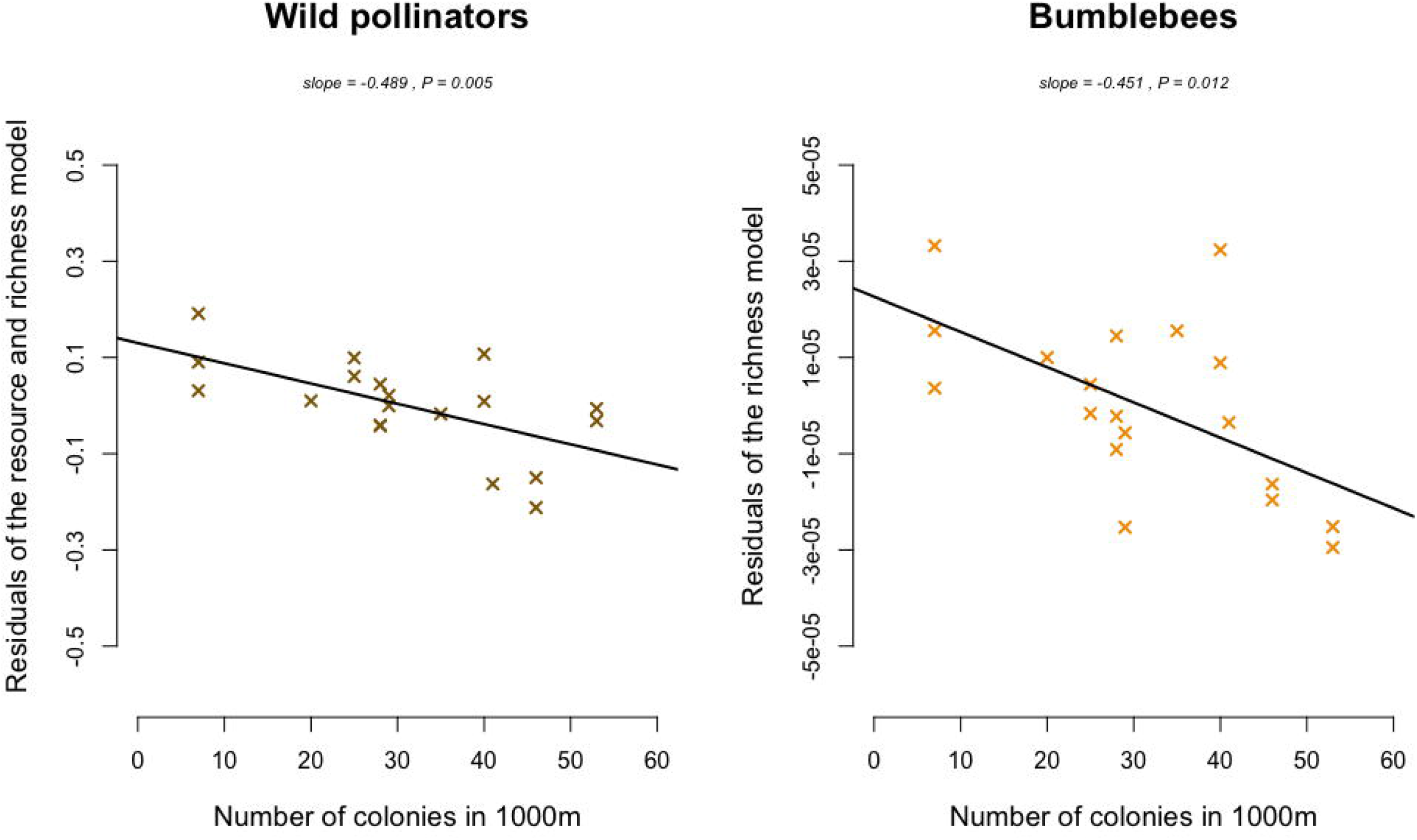
Morphological group visitation rates along the gradient of honey bee colony number in 1000m around our observation sites. When the best model contained also the estimation of floral resources and/or richness as explanatory variable, residuals of morphological group visitation rates explained by resource variable only were used as y values.

**Table 1.** Detailed effects of honey bee colony densities on wild pollinator visitation rates. Results of the best linear mixed-effects models containing the colonies number as response variable and floral resources and/or richness as covariable for each buffer scale. All variables were scaled. Model selection was performed according to AIC criterion.

| Morphological groups and scales | Predictor | Value | Standard deviation | Degree of freedom | t-value | P-value | AICc |
| --- | --- | --- | --- | --- | --- | --- | --- |
| Null model honey bees | Intercept | -0.008 | 0.272 | 16 | -0.030 | 0.977 | 63.50 |
| Honey bees 500m | Intercept | -0.012 | 0.274 | 15 | -0.044 | 0.966 | **61.00** |
|  | Colonies number | 0.501 | 0.193 | 15 | 2.597 | 0.020 |  |
| Null model wild pollinators | Intercept | -0.012 | 0.269 | 16 | -0.045 | 0.964 | 63.50 |
| Wild pollinators 500m | Intercept | -0.088 | 0.418 | 13 | -0.210 | 0.837 | **47.80** |
|  | Colonies number | -0.614 | 0.133 | 13 | -4.628 | 0.001 |  |
|  | Resources | 0.401 | 0.135 | 13 | 2.967 | 0.011 |  |
|  | Richness | 0.659 | 0.144 | 13 | 4.562 | 0.001 |  |
| Wild pollinators 1000m | Intercept | -0.083 | 0.416 | 13 | -0.200 | 0.844 | **56.30** |
|  | Colonies number | -0.489 | 0.143 | 13 | -3.408 | 0.005 |  |
|  | Resources | 0.186 | 0.142 | 13 | 1.316 | 0.211 |  |
|  | Richness | 0.527 | 0.175 | 13 | 3.008 | 0.010 |  |
| Null model large solitary bees | Intercept | -0.085 | 0.417 | 16 | -0.203 | 0.842 | 59.70 |
| Large solitary bees 500m | Intercept | -0.111 | 0.496 | 14 | -0.224 | 0.826 | **51.30** |
|  | Colonies number | -0.425 | 0.133 | 14 | -3.185 | 0.007 |  |
|  | Resources | 0.668 | 0.134 | 14 | 4.992 | 0.000 |  |
| Null model bumblebees | Intercept | 0.017 | 0.267 | 16 | 0.062 | 0.952 | 63.80 |
| Bumblebees 1000m | Intercept | 0.000 | 0.147 | 14 | 0.000 | 1.000 | **52.70** |
|  | Colonies number | -0.451 | 0.155 | 14 | -2.908 | 0.012 |  |
|  | Richness | 0.561 | 0.155 | 14 | 3.616 | 0.003 |  |
| Null model coleoptera | Intercept | -0.001 | 0.273 | 16 | -0.004 | 0.997 | 63.50 |
| Coleoptera 500m | Intercept | -0.012 | 0.285 | 14 | -0.040 | 0.968 | **56.60** |
|  | Colonies number | -0.671 | 0.174 | 14 | -3.854 | 0.002 |  |
|  | Resources | 0.714 | 0.175 | 14 | 4.092 | 0.001 |  |

### Effect of honey bee colony densities on network structure

Regarding the structure of the pollination networks, we found that the evenness of interactions between wild pollinators, excluding honey bee, and plants was negatively related to honey bee colony density within 1000 meter buffers (1000m – slope = −0.487; p = 0.008 – Fig 5, Table 2).

**Fig 5.**
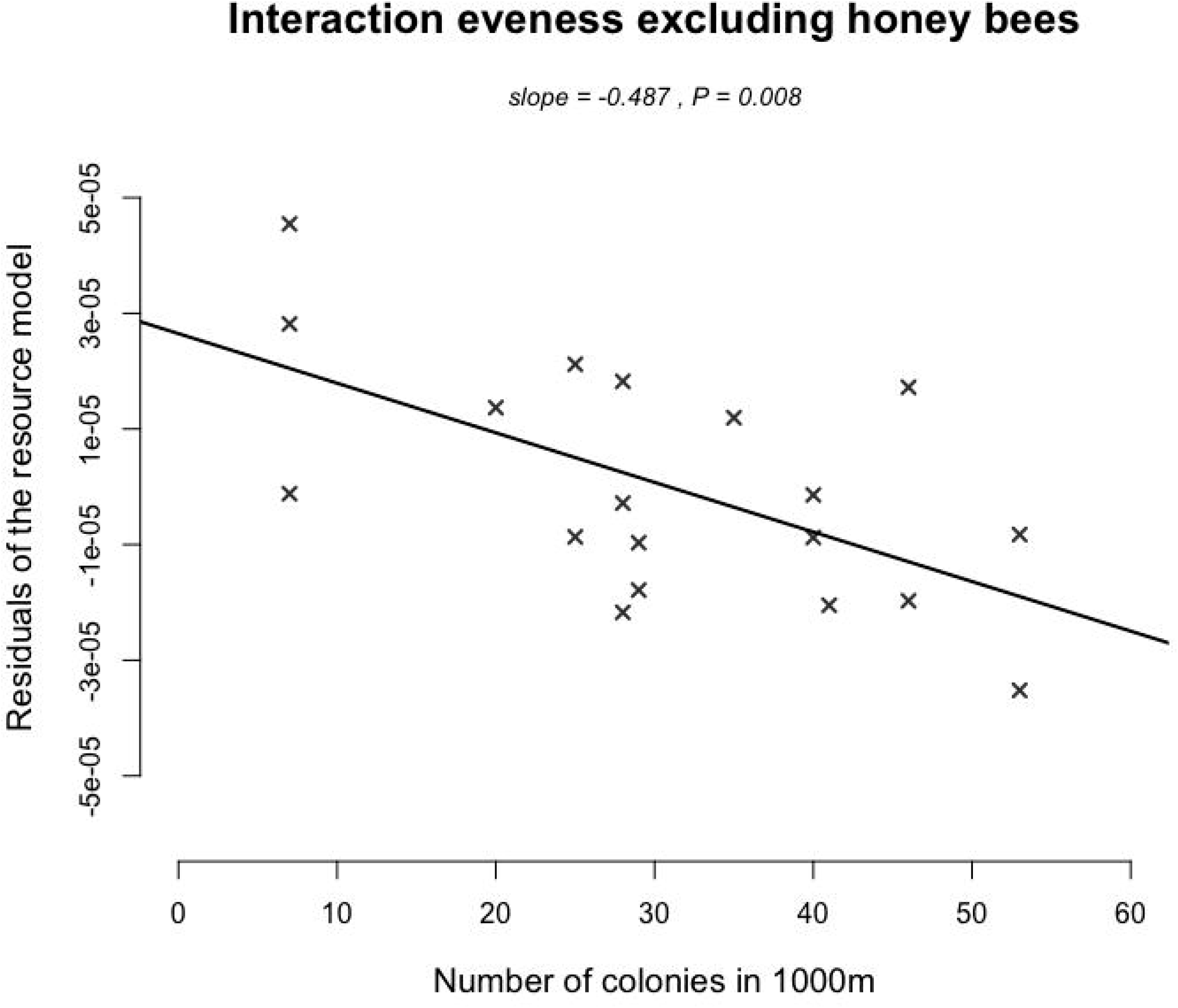
Interaction evenness along the gradient of honey bee colony number in 1000m radius around our observation sites, once the effect of resources was taken into account.

**Table 2.** Detailed effets of honey bee colony densities on the interaction evenness. Results of best linear mixed effects models on interaction eveness containing colonies number and floral resources as covariates. All variables were scaled. Model selection was performed according to AIC criterion.

| Network index | Predictor | Value | Standard deviation | Degree of freedom | t-value | P-value | AICc |
| --- | --- | --- | --- | --- | --- | --- | --- |
| Null model interaction evenness without honey bees | Intercept | 0.000 | 0.229 | 16 | 0.000 | 1.000 | 63.70 |
| Interaction evenness without honey bees 1000m | Intercept | 0.000 | 0.153 | 14 | 0.000 | 1.000 | **54.20** |
|  | Colonies number | -0.487 | 0.159 | 14 | -3.067 | 0.008 |  |
|  | Resources | 0.686 | 0.159 | 14 | 4.320 | 0.001 |  |

### Floral preferences between wild and domesticated pollinators

We found that wild pollinators visited significantly more spontaneous plant species than honey bees (t-test, p = 0.022). Furthermore, honey bees significantly preferred foraging on managed plant species than on spontaneous ones (t-test, p = 0.001; Fig 6) whereas wild pollinators had no preference for a particular plant group, managed and spontaneous plant species being equally visited (t-test, p = 0.745).

**Fig 6.**
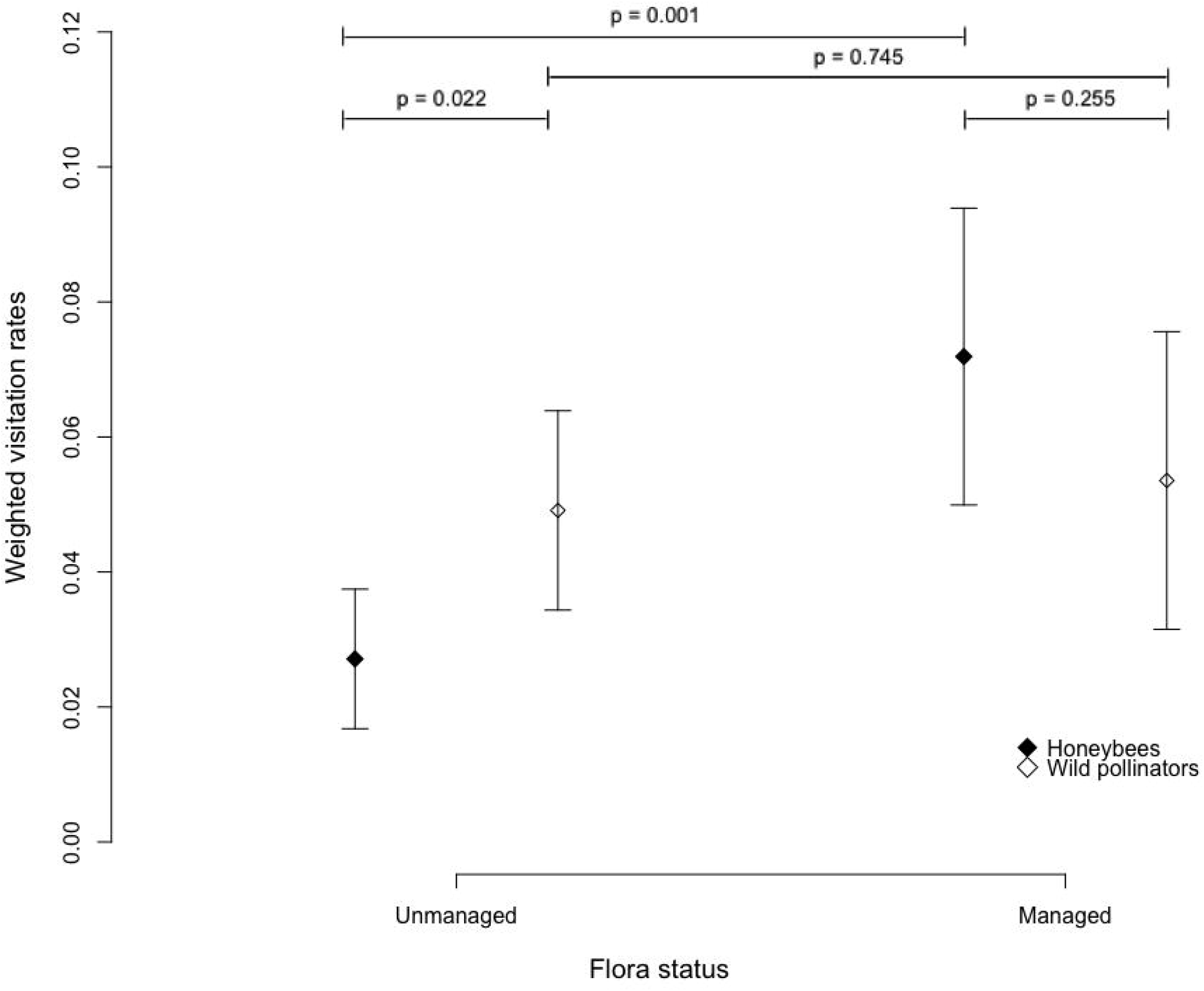
Floral preferences of honey bees and wild pollinators. Mean visitation rate on spontaneous and managed flora weighted by the percentage of managed and spontaneous plant species sampled at each site. Mean and 95% confidence intervals are represented. p-value were obtained with Student’s t-test.

## Discussion

We showed that in the city of Paris, the visitation rate of wild pollinators and especially the pollinating activity of large solitary bees, bumblebees and beetles, was negatively related to the density of honey bee colonies in the surrounding. This first finding resonates with a growing body of literature highlighting a negative effect of high honey bee colony densities on the wild pollinating fauna [11,21]. Although our study is correlative and does not provide direct evidences, our results are consistent with the hypothesis that honey bee might outcompete the wild pollinating fauna by exploiting flowering rewards (nectar and pollen) more efficiently [18,19,33].

The negative correlation between the visitation rates of the total wild fauna and the honey bee colony density was found for both scales, within 500 and 1000 meter buffers. When focusing on each pollinator morphological group, this effect was however scale dependent. The visitation rate of large solitary bees and beetles was negatively correlated to honey bee colony density within 500 meter buffers whereas the visitation rate performed by honey bees increased. The bumblebee visitation rates were negatively correlated to the honey bee colony density within 1000 meter buffers. Those differences might be partly due to the foraging abilities of these groups. The large solitary bees includes numerous species which can forage from few hundred meters to several kilometers from their nest, depending on the species considered and the landscape context [24,34]. Bumblebees in the other hand are known to forage at large scales, up to 2800 meters from their nest [35]. These two morphological groups of Apidae and honey bees have similar dietary requirements, exploiting the same floral resources (pollen and nectar) [17,36]. As summarized in Wojcik et al. 2018 [37], previous studies have found that adding honey bee colonies could negatively affect wild bees and particularly bumblebees especially due to this overlap in resource use. On the contrary, flies, syrphids and butterflies do not exclusively rely on floral resources, especially during their larval life-stage, which might explain the absence of negative interactions with honey bees [38]. Small solitary bees do require pollen and nectar for both larva and adult stages but because body size and mouthpart length are correlated traits [39], small solitary bees might prefer to seek resources on few plant species (shallow flowers, see [23]). Being opposite, larger pollinators such as honey bee and bumblebees, could preferentially forage on plants best adapted to their morphology (deep flowers see [40,41]). In that way, small solitary bees might be less sensitive to the increase in honey bee colony densities. The sharp decline of beetles’ foraging activity with the honey bee colony density within 500 meter buffers is more challenging to explain. There is little literature on floral preferences of beetles. Also, their foraging range seems to be highly variable. As examples, Englund (1993) found that *Cetonia aurata* had a 18m foraging range and Juhel et al. (2017) estimated the foraging range of *Brassicogethes aeneus* up to 1.2km [42,43]. This underlines the difficulty to relate scale dependent ecological effects with ecological traits of species. For honey bees, we did not detect any increase in their visitation rate with honey bee colony density within 1000 meter buffers. Honey bee foraging range seems to be highly context dependent, from several hundred meters to several kilometers [44,45]. Additionally, Couvillon et al. 2015 demonstrated that honey bee foraging distances both depend on the type of rewards that honey bees seek (nectar or pollen) and on the month considered [46]. The scale to which organisms respond to landscape characteristics thus appear dependent of the context and sensitive to various components acting together. In dense urban habitats, pollinator’s foraging distance might also be sensitive to building height, width or to the spatial distribution of green spaces and floral resources [47]. But at this stage, we cannot exclude that the observed decline in the foraging activity of some morphological groups could be linked to another variable not considered in this study. First, some previous studies did not highlight competitive effects of honey bees on other flower-visitors [48] or even the opposite with wild bumblebees negatively affecting honey bee foraging activity [49]. Secondly, most of studies like ours, exploring the potential negative impact of honey bee on the wild fauna only linked the density of colony with the foraging activity of the wild fauna [18,50]. However, as underlined by Mallinger et al., 2017 [21] other critical parameters such as wild pollinator reproduction success (fitness), population or community dynamics are rarely explored which impedes us to draw clear recommendations to land managers. This sheds a light on the importance of conducing new investigations on the links between honey bee colony density and the wild fauna to better understand the underlying mechanisms.

We also recorded a decrease in the evenness of plant-pollinator interaction networks with the honey bee colony densities within 1000 meter buffers. Interaction evenness decreases when the network is dominated by few and/or highly frequent interactions. A high evenness has been previously associated with a good network stability [51,52]. Being opposite, a low interaction evenness has been highlighted in degraded ecosystems [53] and in invaded networks [54]. In a previous meta-analysis [11], we showed that the honey bee position in interaction networks is comparable to that often found for invasive pollinators. Here, the lower evenness at high colonies densities within 1000 meter buffers should be more linked to the decrease of wild pollinators and particularly of bumblebee’s visitation rate. This question the potential impact of urban beekeeping on the whole interaction network and urges once again the need for news studies regarding this topic.

In parallel, we showed that honey bees tended to significantly focus their visits on managed plant species, whereas wild pollinators did not show preferences between managed and spontaneous plants. Honey bees often focus their visits on the most abundant resources to cover the colony needs [55] and ornamental flowerbeds might thus be attractive for them. Among the species most visited by honey bees, we indeed found *Lavendula sp.* and *Geranium sanguineum* which are common in ornamental flowerbeds. In the other hand, spontaneous flowers received significantly more visits from wild pollinators and might rely more on the wild fauna for pollination. The observed decline of the wild fauna visitation rate associated with high colony densities may have negative consequences for the reproduction of this spontaneous flora. Nevertheless, several other factors might explain insect’s flowers preferences and foraging choices such as the morphology, the color, the amount of resources, or the life span of flowers [56].

Numerous cities around the globe have experienced recent and fast increases in honey bee colony densities. The average colony density in Paris is higher than the national level but is far below other cities like Brussels or London (respectively 6.5, 2.5, 15 and 10 colonies/km^2^ [13,14,57]). Cities also harbor a non-negligible diversity of wild pollinators [58–60]. Altogether, our results question the fast development of urban beekeeping and the enthusiasm of citizens and mass media for the installation of hives in cities and the urban management practices supposedly conducted to sustain biodiversity. This underlines the need for new studies exploring how domestic and wild pollinators coexist in urban habitats. In conclusion, we suggest that stakeholders should take into account the impacts that apiaries could have on the wild fauna [19,33]. If the capacity of urban ecosystems to provide the pollination function is to be preserved, land owners may focus their management practices on increasing floral resources and nesting habitats for pollinators in urban environments instead of adding honey bee colonies.

## Supporting Information

S1 Table. List of plant species with their status.

S2 Table. Distance between study sites in meters.

S3 Table. Open floral unit number per m^2^ of vegetative cover at the peak of flowering season from AgriLand Database.

S4 Table. Number of plants status sampled per year.

S5 Table. Results of morphological groups’ model selection based on AIC criterion.

S6 Table. Results of interaction evenness model selection based on AIC criterion.

S1 Data. Dataset of the study.

## Acknowledgements

We thank M. Henry, G. Rodet and M. Baude for contributing to ideas and discussions throughout this project. We also thank E. Meineri for advices about statistics models and spatial autocorrelation tests. We are grateful to F. Flacher, L. Schurr and L. Affre for revising versions of the manuscript. We thank all those who assisted with the fieldwork and volunteer beekeepers involved in the Observatoire Francilien des Abeilles.

## Author contributions

L.R., B.G. and A.M. compiled and analysed data; L.R. B.G. C.F. and I.D. discussed and revised versions of the manuscript.

