## Supplementary material for "Wild pollinator activities negatively related to honey bee colony densities in urban context": S1, S2, S3, S4, S5, S6 Table

**S1 Table. List of plant species with their status.**

| **Species** | **Spontaneous** | **Managed** | **Species** | **Spontaneous** | **Managed** |
| --- | --- | --- | --- | --- | --- |
| *Alcea rosea* L., 1753 | X | X | *Papaver croceum* Ledeeb., 1830 |  | X |
| *Bellis perennis* L., 1753 | X | X | *Passiflora caerulea* L., 1753 |  | X |
| *Potentilla indica* (Andrews) Th.Wolf, 1904 | X | X | *Pelargonium x grandiflorum* |  | X |
| *Sedum album* L., 1753 | X | X | *Perovskia atriplicifolia* Benth., 1848 |  | X |
| *Abelia* x *grandiflora* (Rovelli ex André) Rehder, 1900 | | X | *Petunia x atkinsiana* D.Don, 1839 |  | X |
| *Achillea ‘Coronation gold’* |  | X | *Phacelia tanacetifolia* Benth., 1837 |  | X |
| *Aesculus hippocastanum* L., 1753 |  | X | *Philadelphus ‘lewisii’* |  | X |
| *Agapanthus* sp. |  | X | *Philadelphus coronarius* L., 1753 |  | X |
| *Ailanthus altissima* (Mill.) Swingle, 1916 |  | X | *Phyla nodiflora var. canescens* (L.) Greene, 1899 | | X |
| *Albizia* sp. |  | X | *Pittosporum* sp. |  | X |
| *Alisma lanceolatum* With., 1796 |  | X | *Potentilla recta* L., 1753 |  | X |
| *Allium angulosum* L., 1753 |  | X | *Primula vulgaris* Huds., 1762 |  | X |
| *Allium sativum* L., 1753 |  | X | *Prunus laurocerasus* L., 1753 |  | X |
| *Allium schoenoprasum* L., 1753 |  | X | *Prunus* sp. |  | X |
| *Amelanchier canadensis* (L.) Medik., 1793 |  | X | *Pulmonaria rubra ‘Redstart’* |  | X |
| *Anemone sylvestris* L., 1753 |  | X | *Rosa* sp. |  | X |
| *Angelica archangelica* L., 1753 |  | X | *Rosmarinus officinalis* L., 1753 |  | X |
| *Aquilegia vulgaris* L., 1753 |  | X | *Rudbeckia hirta* L., 1753 |  | X |
| *Aster amellus* L., 1753 |  | X | *Rudbeckia* sp. |  | X |
| *Astilbe* sp. |  | X | *Salix* sp. |  | X |
| *Astrantia major* L., 1753 |  | X | *Salvia grahamii* Benth., 1830 |  | X |
| *Berberis aquifolium* Pursh, 1814 |  | X | *Salvia nemorosa* L., 1762 |  | X |
| *Bidens triplinervia* var. *macrantha* (Wedd.) Sherff, 1925 | | X | *Salvia uliginosa* Benth., 1833 |  | X |
| *Borago officinalis* L., 1753 |  | X | *Saxifraga x arendsii* Arends |  | X |
| *Buddleja davidii* Franch., 1887 |  | X | *Securigera varia* (L.) Lassen, 1989 |  | X |
| *Calendula officinalis* L., 1753 |  | X | *Senecio greyii* Hook.f., 1853 |  | X |
| *Camelia* sp. |  | X | *Senecio leucostachys* Baker |  | X |
| *Campanula isophylla* Moretti |  | X | *Solanum lycopersicum* L., 1753 |  | X |
| *Campanula persicifolia* L., 1753 |  | X | *Spiraea japonica* Siebold, 1826 |  | X |
| *Campanula portenschlagiana* Roem. & Schult., 1819 | | X | *Spiraea* sp. |  | X |
| *Campanula rapunculoides* L., 1753 |  | X | *Spiraea vanhouttei* Carrière, 1876 |  | X |
| *Campanula trachelium* L., 1753 |  | X | *Staphisagria macrosperma* Spach, 1838 |  | X |
| *Centaurea benedicta* (L.) L., 1763 |  | X | *Symphytum tuberosum* L., 1753 |  | X |
| *Centaurea gymnocarpa* Moris & De Not. |  | X | *Tagetes patula* L., 1753 |  | X |
| *Centaurea jacea* L., 1753 |  | X | *Tagetes* sp. |  | X |
| *Centranthus ruber* (L.) DC., 1805 |  | X | *Tamarix* sp. |  | X |
| *Ceratostigma plumbaginoides* Bunge, 1833 |  | X | *Teucrium fruticans* L., 1753 |  | X |
| *Chaenomeles japonica* (Thunb.) Lindl. ex Spach, 1834 | | X | *Ulex europaeus* L., 1753 |  | X |
| *Choisya ternata* Kunth, 1823 |  | X | *Verbena bonariensis* L., 1753 |  | X |
| *Cistus* sp. |  | X | *Viburnum davidii* Franch., 1886 |  | X |
| *Clinopodium nepeta* (L.) Kuntze, 1891 |  | X | *Viburnum lantana* L., 1753 |  | X |
| *Coreopsis grandiflora* Hogg ex Sweet, 1836 |  | X | *Viburnum plicatum* Thunb., 1784 |  | X |
| *Cornus alba* L., 1767 |  | X | *Viburnum* sp. |  | X |
| *Cornus sanguinea* L., 1753 |  | X | *Vinca minor* L., 1753 |  | X |
| *Cosmos bipinnatus* Cav., 1791 |  | X | *Viola cornuta* L., 1763 |  | X |
| *Cotinus x ‘grace’* |  | X | *Zinnia haageana* Regel, 1861 |  | X |
| *Cotoneaster franchetii* Bois, 1902 |  | X | *Achillea millefolium* L., 1753 | X |  |
| *Cotoneaster* sp. |  | X | *Ajuga reptans* L., 1753 | X |  |
| *Cyanus montanus* (L.) Hill, 1768 |  | X | *Alliaria petiolata* (M.Bieb.) Cavara & Grande, 1913 | X |  |
| *Cytisus scoparius* (L.) Link, 1822 |  | X | *Althaea officinalis* L., 1753 | X |  |
| *Dasiphora fruticosa* (L.) Rydb., 1898 |  | X | *Angelica sylvestris* L., 1753 | X |  |
| *Delphinium consolida* L., 1753 |  | X | *Bryonia dioica* Jacq., 1774 | X |  |
| *Delphinium* sp. |  | X | *Convolvulus sepium* L., 1753 | X |  |
| *Deutzia gracilis* Siebold & Zucc., 1839 |  | X | *Centaurium erythraea* Rafn, 1800 | X |  |
| *Deutzia* sp. |  | X | *Cerastium fontanum* Baumg., 1816 | X |  |
| *Deutzia x elegantissima* |  | X | *Chelidonium majus* L., 1753 | X |  |
| *Dianthus barbatus* L., 1753 |  | X | *Cirsium arvense* (L.) Scop., 1772 | X |  |
| *Dianthus carthusianorum* L., 1753 |  | X | *Convolvulus arvensis* L., 1753 | X |  |
| *Dianthus caryophyllus* L., 1753 |  | X | *Crataegus monogyna* Jacq., 1775 | X |  |
| *Dianthus* sp. |  | X | *Crepis capillaris* (L.) Wallr., 1840 | X |  |
| *Diascia* sp. |  | X | *Crepis setosa* Haller f., 1797 | X |  |
| *Digitalis purpurea* L., 1753 |  | X | *Daucus carota* L., 1753 | X |  |
| *Dipelta floribunda* |  | X | *Dipsacus fullonum* L., 1753 | X |  |
| *Doronicum orientale* Hoffm., 1808 |  | X | *Echium vulgare* L., 1753 | X |  |
| *Dorycnium pentaphyllum* Scop., 1772 |  | X | *Epilobium angustifolium* L., 1753 | X |  |
| *Ecballium elaterium* (L.) A.Rich., 1824 |  | X | *Epilobium hirsutum* L., 1753 | X |  |
| *Echinops ritro* L., 1753 |  | X | *Epilobium montanum* L., 1753 | X |  |
| *Erigeron annuus* (L.) Desf., 1804 |  | X | *Epilobium tetragonum* L., 1753 | X |  |
| *Eruca sativa* Mill., 1768 |  | X | *Geranium molle* L., 1753 | X |  |
| *Erysimum cheiri* (L.) Crantz, 1769 |  | X | *Geranium robertianum* L., 1753 | X |  |
| *Eschscholzia californica* Cham., 1820 |  | X | *Glechoma hederacea* L., 1753 | X |  |
| *Euphorbia amygdaloides* subsp. *robbiae* (Turrill) Stace, 1989 | | X | *Hyacinthoides non-scripta* (L.) Chouard ex Rothm., 1944 | X |  |
| *Euphorbia characias* L., 1753 |  | X | *Hypericum perforatum L., 1753* | X |  |
| *Forsythia* sp. |  | X | *Hypochaeris radicata* L., 1753 | X |  |
| *Fothergilla ‘major’* |  | X | *Jacobaea vulgaris* Gaertn., 1791 | X |  |
| *Fuchsia magellanica* Lam., 1788 |  | X | *Lactuca muralis* (L.) Gaertn., 1791 | X |  |
| *Geranium phaeum* L., 1753 |  | X | *Lamium album* L., 1753 | X |  |
| *Geranium sanguineum* L., 1753 |  | X | *Lamium purpureum* L., 1753 | X |  |
| *Gerbera aurantiaca* |  | X | *Lathyrus latifolius* L., 1753 | X |  |
| *Glycyrrhiza glabra* L., 1753 |  | X | *Leucanthemum vulgare* Lam., 1779 | X |  |
| *Hebe colensoi glauca* |  | X | *Lotus corniculatus* L., 1753 | X |  |
| *Helianthemum grandiflorum* (Scop.) DC., 1805 | | X | *Lysimachia arvensis* (L.) U.Manns & Anderb., 2009 | X |  |
| *Helianthemum nummularium* (L.) Mill., 1768 |  | X | *Lythrum salicaria* L., 1753 | X |  |
| *Helianthemum* sp. |  | *X* | *Malva neglecta* Wallr., 1824 | X |  |
| *Helianthemum violaceum* (Cav.) Pers., 1806 |  | X | *Malva sylvestris* L., 1753 | X |  |
| *Hibiscus palustris* L., 1753 |  | X | *Matricaria inodora* Lam., 1779 | X |  |
| *Hibiscus trionum* L., 1753 |  | X | *Matricaria recutita* L., 1753 | X |  |
| *Hosta plantaginea* |  | X | *Medicago lupulina* L., 1753 | X |  |
| *Houttuynia cordata* |  | X | *Melilotus albus* Medik., 1787 | X |  |
| *Hydrangea quercifolia* |  | X | *Melissa officinalis* L., 1753 | X |  |
| *Hydrangea* sp. |  | X | *Mentha aquatica* L., 1753 | X |  |
| *Hylotelephium telephium* (L.) H.Ohba, 1977 |  | X | *Mentha arvensis* L., 1753 | X |  |
| *Hypericum ‘hidcote’* |  | X | *Origanum vulgare* L., 1753 | X |  |
| *Hypericum calycinum* L., 1767 |  | X | *Papaver rhoeas* L., 1753 | X |  |
| *Hypericum lanceolatum* Lam., 1797 |  | X | *Helminthotheca echioides* (L.) Holub, 1973 | X |  |
| *Hypericum* x *moserianum* Luquet ex André, 1888 | | X | *Picris hieracioides* L., 1753 | X |  |
| *Iberis umbellata* L., 1753 |  | X | *Plantago lanceolata* L., 1753 | X |  |
| *Inula spiraeifolia* L., 1759 |  | X | *Plantago major* L., 1753 | X |  |
| *Kerria japonica* (L.) DC., 1818 |  | X | *Poa annua* L., 1753 | X |  |
| *Kniphofia* sp. |  | X | *Potentilla reptans* L., 1753 | X |  |
| *Kolwitzia* sp. |  | X | *Prunella vulgaris* L., 1753 | X |  |
| *Lamium galeobdolon* (L.) L., 1759 |  | X | *Pulicaria dysenterica* (L.) Bernh., 1800 | X |  |
| *Laserpitium gallicum* L., 1753 |  | X | *Ranunculus acris* L., 1753 | X |  |
| *Lavandula angustifolia* Mill., 1768 |  | X | *Ranunculus arvensis* L., 1753 | X |  |
| *Lavandula latifolia* Medik., 1784 |  | X | *Ranunculus repens* L., 1753 | X |  |
| *Lavandula* sp. |  | X | *Rosa rubiginosa* L., 1771 | X |  |
| *Ligustrum lucidum* W.T.Aiton, 1810 |  | X | *Rubus fruticosus* L., 1753 | X |  |
| *Ligustrum ovalifolium* Hassk., 1844 |  | X | *Sambucus nigra* L., 1753 | X |  |
| *Ligustrum vulgare* L., 1753 |  | X | *Saxifraga tridactylites* L., 1753 | X |  |
| *Linum usitatissimum* subsp. *angustifolium* (Huds.) Thell., 1912 | | X | *Senecio inaequidens* DC., 1838 | X |  |
| *Lonicera implexa* Aiton, 1789 |  | X | *Senecio vulgaris* L., 1753 | X |  |
| *Lonicera nitida* E.H.Wilson, 1911 |  | X | *Sisymbrium irio* L., 1753 | X |  |
| *Lonicera periclymenum* L., 1753 |  | X | *Sonchus oleraceus* L., 1753 | X |  |
| *Lunaria annua* L., 1753 |  | X | *Stachys recta* L., 1767 | X |  |
| *Luzula nivea* (Nathh.) DC., 1805 |  | X | *Tanacetum vulgare* L., 1753 | X |  |
| *Magnolia grandiflora* L., 1759 |  | X | *Taraxacum* section *ruderalia* | X |  |
| *Malus* sp. |  | X | *Thymus serpyllum* L., 1753 | X |  |
| *Mantisalca salmantica* (L.) Briq. & Cavill., 1930 | | X | *Thymus vulgare* L., 1753 | X |  |
| *Medicago arborea* L., 1753 |  | X | *Trifolium pratense* L., 1753 | X |  |
| *Mentha spicata* L., 1753 |  | X | *Trifolium repens* L., 1753 | X |  |
| *Monarda* sp. |  | X | *Valerianella locusta* (L.) Laterr., 1821 | X |  |
| *Muscari* sp. |  | X | *Verbascum thapsus* L., 1753 | X |  |
| *Myosotis* sp. |  | X | *Verbena officinalis* L., 1753 | X |  |
| *Narcissus pseudonarcissus* L., 1753 |  | X | *Veronica persica* Poir., 1808 | X |  |
| *Narcissus tazetta* L., 1753 |  | X | *Vicia cracca* L., 1753 | X |  |
| *Nepeta* sp. |  | X | *Vicia sativa* L., 1753 | X |  |
| *Nepeta* x *faassenii* Bergmans ex Stearn, 1950 |  | X | *Vicia sepium* L., 1753 | X |  |
| *Nigella damascena* L., 1753 |  | X | *Viola reichenbachiana* Jord. ex Boreau, 1857 | X |  |
| *Oenothera lindheimeri* (Engelm. & A.Gray) W.L.Wagner & Hoch, 2007 | | X | *Viola* sp. | X |  |

**S2 Table. Distance between study sites in meters.**

| Distance (m) | Site 1 | Site 2 | Site 3 | Site 4 | Site 5 | Site 6 |
| --- | --- | --- | --- | --- | --- | --- |
| Site 2 | 723.84 |  |  |  |  |  |
| Site 3 | 4238.60 | 3732.95 |  |  |  |  |
| Site 4 | 3066.09 | 2559.29 | 1176.64 |  |  |  |
| Site 5 | 5965.41 | 5459.33 | 1729.15 | 2905.76 |  |  |
| Site 6 | 2954.27 | 2528.07 | 1335.80 | 410.04 | 3032.67 |  |
| Site 7 | 881.36 | 846.97 | 4545.26 | 3379.50 | 6264.35 | 3369.72 |

**S3 Table. Open floral unit number per m² of vegetative cover at the peak of flowering season from AgriLand Database.**

| Strata | Floral density at the flowering peak |
| --- | --- |
| Small (max height <1 m) | 2713.6 |
| Medium (1 m ≤ max height <10 m) | 2186.4 |
| Tall (max height ≥ 10 m) | 5291.4 |

**S4 Table. Number of plants status sampled per year.**

| **Year** | **Number of spontaneous plant occurencies sampled for all observation round** | **Number of spontaneous plant occurencies sampled for all observation round** |
| --- | --- | --- |
| 2014 | 106 | 72 |
| 2015 | 148 | 218 |
| 2016 | 116 | 119 |
| Total | 370 | 409 |

**S5 Table. Results of morphological groups’ model selection based on AIC criterion.**

| **Models** |  | **Intercept** | **Colonies** | **Resources** | **Mean Richness** | **Df** | **log Likelihood** | **AICc** | **Delta** | **Weight** |
| --- | --- | --- | --- | --- | --- | --- | --- | --- | --- | --- |
| Honey bees 500m | 2 | -0.012 | 0.501 |  |  | 5 | -23.207 | 61.00 | 0.00 | 0.544 |
|  | 1 | -0.008 |  |  |  | 4 | -26.311 | 63.50 | 2.45 | 0.160 |
| Honey bees 1000m | 1 | 3.878 |  |  |  | 4 | -15.305 | 41.50 | 0.00 | 0.546 |
| Wild pollinators 500m | 8 | -0.088 | -0.614 | 0.401 | 0.659 | 7 | -14.933 | 54.00 | 0.00 | 0.782 |
|  | 6 | -0.079 | -0.414 |  | 0.710 | 6 | -19.126 | 57.30 | 3.20 | 0.158 |
|  | 5 | -0.062 |  |  | 0.634 | 5 | -22.816 | 60.20 | 6.20 | 0.035 |
|  | 4 | -0.033 | -0.622 | 0.497 |  | 6 | -22.138 | 63.30 | 9.23 | 0.008 |
|  | 1 | -0.012 |  |  |  | 4 | -26.341 | 63.50 | 9.49 | 0.007 |
| Wild pollinators 1000m | 6 | -0.084 | -0.450 | 0.581 |  | 6 | -18.645 | 56.30 | 0.00 | 0.644 |
|  | 8 | -0.083 | -0.489 | 0.186 | 0.527 | 7 | -17.611 | 59.40 | 3.11 | 0.136 |
|  | 5 | -0.062 |  |  | 0.634 | 5 | -22.816 | 60.20 | 3.96 | 0.089 |
|  | 2 | -0.038 | -0.522 |  |  | 5 | -23.104 | 60.80 | 4.53 | 0.067 |
|  | 4 | -0.043 | -0.573 | 0.292 |  | 6 | -21.644 | 62.30 | 6.00 | 0.032 |
|  | 1 | -0.012 |  |  |  | 4 | -26.341 | 63.50 | 7.25 | 0.017 |
| Small solitary bees 500m | 2 | -0.011 | -0.400 |  |  | 5 | -24.125 | 62.90 | 0.00 | 0.375 |
|  | 1 | -0.008 |  |  |  | 4 | -26.089 | 63.00 | 0.17 | 0.345 |
| Small solitary bees 1000m | 1 | -0.008 |  |  |  | 4 | -26.089 | 63.00 | 0.00 | 0.347 |
| Large solitary bees 500m | 4 | -0.111 | -0.425 | 0.668 |  | 6 | -16.163 | 51.30 | 0.00 | 0.753 |
|  | 8 | -0.125 | -0.422 | 0.641 | 0.178 | 7 | -15.278 | 54.70 | 3.41 | 0.137 |
|  | 3 | -0.101 |  | 0.450 |  | 5 | -20.594 | 55.80 | 4.48 | 0.08 |
|  | 7 | -0.114 |  | 0.427 | 0.164 | 6 | -20.188 | 59.40 | 8.05 | 0.013 |
|  | 1 | -0.085 |  |  |  | 4 | -24.444 | 59.70 | 8.42 | 0.011 |
| Large solitary bees 1000m | 3 | -0.094 |  | 0.466 |  | 5 | -20.173 | 55.00 | 0.00 | 0.498 |
|  | 4 | -0.106 | -0.256 | 0.505 |  | 6 | -18.391 | 55.80 | 0.82 | 0.33 |
|  | 7 | -0.106 |  | 0.442 | 0.138 | 6 | -19.882 | 58.80 | 3.80 | 0.074 |
|  | 1 | -0.085 |  |  |  | 4 | -24.444 | 59.70 | 4.78 | 0.046 |
| Bumblebees 500m | 5 | 0.000 |  |  | 0.665 | 5 | -20.901 | 56.40 | 0.00 | 0.674 |
|  | 7 | 0.000 |  | 0.228 | 0.622 | 6 | -20.008 | 59.00 | 2.60 | 0.184 |
|  | 6 | 0.000 | -0.059 |  | 0.668 | 6 | -20.84 | 60.70 | 4.26 | 0.08 |
|  | 8 | 0.000 | -0.229 | 0.344 | 0.611 | 7 | -19.245 | 62.70 | 6.25 | 0.03 |
|  | 1 | 0.017 |  |  |  | 4 | -26.363 | 63.60 | 7.17 | 0.019 |
| Bumblebees 1000m | 6 | 0.000 | -0.451 |  | 0.561 | 6 | -16.87 | 52.70 | 0.00 | 0.614 |
|  | 8 | 0.000 | -0.502 | 0.247 | 0.506 | 7 | -15.267 | 54.70 | 1.98 | 0.229 |
|  | 5 | 0.000 |  |  | 0.665 | 5 | -20.901 | 56.40 | 3.68 | 0.098 |
|  | 2 | 0.012 | -0.565 |  |  | 5 | -22.448 | 59.50 | 6.77 | 0.021 |
|  | 4 | 0.017 | -0.612 | 0.353 |  | 6 | -20.323 | 59.60 | 6.91 | 0.019 |
|  | 7 | 0.000 |  | 0.147 | 0.639 | 6 | -20.537 | 60.10 | 7.33 | 0.016 |
|  | 1 | 0.017 |  |  |  | 4 | -26.363 | 63.60 | 10.84 | 0.003 |
| Coleoptera 500m | 4 | -0.012 | -0.671 | 0.714 |  | 6 | -18.778 | 56.60 | 0.00 | 0.83 |
|  | 8 | -0.018 | -0.665 | 0.686 | 0.154 | 7 | -18.352 | 60.90 | 4.33 | 0.095 |
|  | 1 | -0.001 |  |  |  | 4 | -26.299 | 63.50 | 6.90 | 0.026 |
| Coleoptera 1000m | 1 | -0.001 |  |  |  | 4 | -26.299 | 63.50 | 0.00 | 0.432 |
| Lepidoptera 500m | 1 | -0.010 |  |  |  | 4 | -26.392 | 63.60 | 0.00 | 0.626 |
| Lepidoptera 1000m | 1 | -0.010 |  |  |  | 4 | -26.392 | 63.60 | 0.00 | 0.565 |
| Syrphids 500m | 1 | -0.041 |  |  |  | 4 | -26.001 | 62.90 | 0.00 | 0.506 |
| Syrphids 1000m | 1 | -0.041 |  |  |  | 4 | -26.001 | 62.90 | 0.00 | 0.38 |
| Diptera 500m | 5 | -0.077 |  |  | 0.664 | 5 | -24.289 | 63.20 | 0.00 | 0.404 |
|  | 1 | 0.000 |  |  |  | 4 | -26.446 | 63.70 | 0.56 | 0.306 |
| Diptera 1000m | 5 | -0.077 |  |  | 0.664 | 5 | -24.289 | 63.20 | 0.00 | 0.445 |
|  | 1 | 0.000 |  |  |  | 4 | -26.446 | 63.70 | 0.56 | 0.337 |

**S6 Table. Results of interaction evenness model selection based on AIC criterion.**

| **Models** |  | **Intercept** | **Colonies** | **Resources** | **Mean Richness** | **Df** | **log Likelihood** | **AICc** | **Delta** | **Weight** |
| --- | --- | --- | --- | --- | --- | --- | --- | --- | --- | --- |
| Interaction evenness without honey bees 500m | 3 | 0.000 |  | 0.590 |  | 5 | -22.375 | 59.40 | 0.00 | 0.663 |
|  | 4 | 0.000 | -0.238 | 0.709 |  | 6 | -21.735 | 62.50 | 3.10 | 0.141 |
|  | 7 | 0.000 |  | 0.576 | 0.076 | 6 | -22.294 | 63.60 | 4.22 | 0.080 |
|  | 1 | 0.000 |  |  |  | 4 | -26.446 | 63.70 | 4.38 | 0.074 |
| Interaction evenness without honey bees 1000m | 4 | 0.000 | -0.487 | 0.686 |  | 6 | -17.599 | 54.20 | 0.00 | 0.819 |
|  | 3 | 0.000 |  | 0.612 |  | 5 | -21.993 | 58.60 | 4.40 | 0.091 |
|  | 8 | 0.000 | -0.501 | 0.698 | -0.054 | 7 | -17.537 | 59.30 | 5.06 | 0.065 |
|  | 7 | 0.000 |  | 0.598 | 0.079 | 6 | -21.900 | 62.80 | 8.60 | 0.011 |
|  | 1 | 0.000 |  |  |  | 4 | -26.446 | 63.70 | 9.55 | 0.007 |
